# miR-24:Prdx6 interactions regulate oxidative stress and viability of myogenic progenitors during ageing

**DOI:** 10.1101/2021.01.25.428069

**Authors:** Ana Soriano-Arroquia, John Gostage, David Bardell, Eugene McCloskey, Ilaria Bellantuono, Peter Clegg, Brian McDonagh, Katarzyna Goljanek-Whysall

**Affiliations:** Institute of Life Course and Medical Sciences, University of Liverpool William Henry Duncan Building. 6 West Derby Street, Liverpool L7 8TX, UK; The Medical Research Council / Versus Arthritis Centre for Integrated Research into Musculoskeletal Ageing; CIMA. University of Liverpool. William Henry Duncan Building. 6 West Derby Street, Liverpool L7 8TX, UK; Discipline of Physiology, School of Medicine, National University of Ireland Galway, Ireland; Healthy Lifespan Institute and The Centre for Integrated Research in Musculoskeletal Ageing, Department of Oncology and Metabolism, University of Sheffield, Sheffield, UK

**Author notes:** Correspondence: Katarzyna Goljanek-Whysall; Institute of Life Course and Medical Sciences, University of Liverpool and Discipline of Physiology, School of Medicine, National University of Ireland Galway, Ireland.

**Keywords:** microRNA, muscle regeneration, ageing, satellite cells, sarcopenia, Prdx6, miR-24, reactive oxygen species, ROS

## Abstract

microRNAs regulate a myriad of physiological processes, including skeletal muscle regeneration and homeostasis. During ageing, changes in muscle fibre microenvironment contribute to the capability of satellite cells to regenerate the muscle in response to injury and loading stressors. In this study, we isolated murine satellite cells and primary myogenic progenitors from mice and humans to demonstrate that the microRNA miR-24-3p and its target peroxiredoxin 6 (Prdx6) play an important role in muscle regeneration during ageing, regulating satellite cell viability and their differentiation potential. Our results show upregulation of miR-24 during early stages of muscle regeneration *in vivo* in adult mice, suggesting a potential role of miR-24 at the early stages of muscle injury. On contrary, miR-24 was downregulated during regeneration of muscle of old mice. miR-24 was also downregulated, whereas its target gene Prdx6 was upregulated, in satellite cells isolated from old mice. miR-24 consistently regulated viability and myogenic potential of myogenic progenitors from both humans and old mice, suggesting that changes in miR-24 levels during ageing may contribute to defective early stages of muscle regeneration during ageing through affecting satellite cell viability and myogenic potential. This regulation likely occurs *via* miR-24 counteracting the generation of reactive oxygen species through Prdx6 de-repression in primary myogenic progenitors isolated from humans and old mice. We propose that downregulation of miR-24 in muscle of old mice following injury may be a protective mechanism against elevated ROS levels to maintain satellite cell viability and myogenic potential, acting through Prdx6 upregulation. However, as miR-24 is a regulator of p16 and p21, this downregulation may lead to increased satellite cell senescence, therefore representing an age-related failed compensatory mechanism.

## Introduction

The regenerative capacity of skeletal muscle facilitates a high plasticity for adaptation to diverse metabolic conditions and energetic demands. Skeletal muscle regeneration after injury and loading stressors relies on satellite cells, the adult muscle stem cells able to regenerate muscle fibres *in vivo*. Following injury and in response to stress signals, satellite cells re-enter the cell cycle, proliferate and either return back to quiescence to maintain the stem cell pool or differentiate and fuse to the existing muscle fibres in order to repair the damage [1]. A balance between self-renewal and myogenic differentiation is essential for successful muscle regeneration after injury [2]. However, the effectiveness of muscle regeneration throughout lifespan not only relies on the functionality of satellite cells [3–5], but also other factors, such as disrupted intracellular signalling and an altered muscle fibre microenvironment, are known to play a key role in muscle wasting during disuse, ageing and chronic diseases [4, 6–9]. In particular, oxidative stress has been demonstrated to alter the cellular microenvironment, resulting in disrupted cellular signalling and potentially oxidative modifications of muscle contractile proteins [10, 11]. The imbalance in myofibrillar redox environment during ageing can result in an accumulation of ROS that contributes to disrupted signalling and function of satellite cells, cellular apoptosis, muscle atrophy and premature ageing [12–14]. Some of the important muscle antioxidant proteins that directly affect intracellular ROS concentrations are members of the peroxiredoxin family: (PRDX1-PRDX6). Peroxiredoxins have the capacity to regulate redox homeostasis and signalling pathways involved in processes such as apoptosis and cell survival or in response to injury. Particularly, Peroxiredoxin 6 (Prdx6) has been demonstrated to regulate both myogenesis and adipogenesis *via* the control of glucose uptake [15, 16], and *Prdx6*^−/−^ mice display increased levels of markers of senescence, metabolic sarcopenia and loss of muscle strength [17].

microRNAs (miRNAs, miRs) are short non-coding RNAs approximately 20-22 nucleotides in length. miRs show partial complementarity to their target mRNA(s) and regulate gene expression at the post-transcriptional level [18, 19]. miRs are known to regulate a myriad of biological processes, including muscle homeostasis and ageing through processes such as ROS generation and scavenging [20–24]. Several studies also demonstrated the regulation of redox balance in muscle by miRs [10]. However, little is known about changes in miR expression in satellite cells during ageing and their functional consequences on muscle homeostasis. Interestingly, miR-24 and the cluster from which it is produced, miR-23–27–24, are highly expressed in skeletal muscle [25]. miR-24 has been proposed to regulate myogenesis *in vitro* and to inhibit muscle fibrosis *in vivo* [26, 27]. miR-24 has been also shown to regulate the expression of tumour suppressor/senescence-associated proteins in a different manner depending on the cell type and metabolic state of the cells [28–31], and regulate cell proliferation, migration, apoptosis and senescence in several tissues and cell types. Yet, the functional role of miR-24 in human and mouse primary myogenic stem/progenitor cells, including oxidative stress, and in skeletal muscle ageing remains elusive.

In this study, we identified the changes in miR-24:Prdx6 interactions in satellite cells during ageing. Our data confirm a decline in satellite cell fraction, viability and myogenic potential in muscle from old mice. miR-24 expression was downregulated in Fluorescence Activated Cell Sorting (FACS)-sorted satellite cells during ageing, with the concomitant upregulation of its target Prdx6. Our results demonstrate an early upregulation of miR-24 in regenerating muscle from adult mice after acute injury which is suppressed in old mice. Using GFP reporter constructs, we demonstrated that miR-24 directly regulated the expression of Prdx6. Changes in miR-24:Prdx6 interactions were associated with altered ROS generation in myogenic progenitors and affected their viability. The effects of miR-24 up- and downregulation were more pronounced in myogenic progenitors from old mice, suggesting a context-dependent role of miR-24 in these cells. We propose that changes in miR-24:Prdx6 interactions during ageing are aimed at preserving satellite cells viability and their ability to regenerate injured myofibres. As the acute increase in ROS following injury is required for effective muscle regeneration, we hypothesise that age-related downregulation of miR-24 and subsequent increased Prdx6 expression in quiescent satellite cells represents a compensatory mechanism against a disrupted redox environment and consequently impaired regenerative capacity of skeletal muscle.

## Materials and Methods

### Mouse samples

All experiments described herein received the ethical approval from The University of Liverpool Animal Welfare and Ethical Review Body (AWERB) and were performed in accordance with UK Home Office guidelines under the UK Animals (Scientific Procedures) Act 1986. All mice were male wild-type C57Bl/6 from Charles River (Margaret), maintained under specific-pathogen free conditions and fed *ad libitum* and maintained under barrier on a 12 hours light/dark cycle. For muscle regeneration, tibialis anterior muscle was injured by intramuscular injection of barium chloride (1.2% in saline). Tissue was collected 1, 7, 14, or 21 days after injury. Muscle was snap-frozen in liquid nitrogen and stored at −80°C as previously described [32]. Primary muscle progenitor cells and satellite cells were directly isolated from fresh lower limbs muscles (extensor digitorum longus, tibialis anterior, gastrocnemius, quadriceps and soleus). For each experiment, n=3-7 biological replicates per group were used. Young: 6-12 weeks old; adult: 6-8 months old; old: 20-24 months old. For miR-24 and Prdx6 expression in FACS-sorted satellite cells: adult: 1-8 months old; old: 20-24 months old.

### Human samples

All experiments described herein involving human samples were performed according to good practice guidance and in accordance with The University of Liverpool, University Hospital Aintree Hospital and South West Wales Research Ethics Committee (Approval No: 13/WA/0374). The University of Liverpool acted as the ethics sponsor for this study. All the donors had given informed consent for enrolment in this study. Muscle biopsies were obtained from foot surgeries (extensor digitorum brevis, tibialis anterior or abductor hallucis muscles) of female patients treated for Hallux Valgus, with an average age of 33 ± 6.78 years old and a Body Mass Index (BMI) < 25. For each experiment, and due to limitations in sample availability, both human primary myogenic progenitors isolated from female donors (n=2-5 per experiment) and commercialised human primary skeletal muscle progenitors (ThermoFisher Scientific, Cat. Number: A12555, n=1-2 per experiment) were used. For all the experiments n=3-7 biological replicates per group, unless stated otherwise.

### Satellite cell isolation

Satellite cells were isolated using FACS as previously described [32, 33]. Briefly, skeletal muscle was isolated from the hind limbs of C57Bl/6 wild type male mice and enzymatically digested with 1.5 U mL-1 collagenase D, 2.4 U mL-1 dispase II and 2.5 mM CaCl2. Cells were then dissolved in sterile FACS buffer (2% horse serum in DPBS), filtered through a 40μm cell strainer and stained with conjugated antibodies in the dark for 30 minutes on ice. Doublets were discriminated and haematopoietic and endothelial cells (PE-CD31^+^/CD45^+^) were excluded from the sorting gates. Satellite cell population was isolated as BV421-CD34^+^, Alexa647-Alpha7Integrin^+^, FICT-Sca1^−^, PE-CD31^−^, PE-CD45^−^ and eFluor780-Viability^−^ dye. Sorting was performed at 4°C and samples were collected in growth media (high-glucose DMEM supplemented with 10% FBS, 1% L-glutamine and 1% penicillin/streptomycin). Sorted cells were immediately centrifuged and resuspended in Qiazol (Qiagen, Cat# 217004) for total RNA isolation.

### RNA isolation

For RNA isolation, cells were collected 48 hr after transfection. Total RNA from sorted cells was isolated using miRNeasy Mini Kit (Qiagen, Cat# 217004). Total RNA from human and mouse primary muscle progenitor cells was isolated using TRIzol/Chloroform standard protocol. After isolation, if necessary samples were purified using ethanol and sodium acetate. RNA concentration and quality were assessed using Nanodrop 2000.

### Real-Time qPCR

cDNA synthesis and Real-Time qPCR were performed as previously described [32]. Briefly, cDNA synthesis was performed from 500ng of RNA (for mRNA) or 100ng of RNA (for microRNA) using SuperScript II (ThermoFisher, Ref. 18064014) or miRscript RT kit II (Qiagen, Ref. 218161), respectively. SYBR Green Mastermix (Qiagen, Ref. 218073) or SsoAdvanced Universal SYBR Green Supermix (BioRad, Ref. 1725271) were used for Real-Time quantitative PCR. Relative expression to β-actin, 18S, β −2 microglobulin (mRNA) or Snord-61 (microRNA) was calculated using delta C_t_ method [32].

### Isolation of primary muscle progenitor cells from mouse and human skeletal muscles

The isolation of human and mouse primary muscle progenitor cells was performed as previously described [34]. Briefly, skeletal muscle tissue was enzymatically digested with 1.5 U mL^−1^ collagenase D (Roche, Ref. 11088882001), 2.4 U mL^−1^ dispase II (Sigma-Aldrich, Ref. D4693) and 2.5 mM CaCl2 (Sigma-Aldrich, Ref. 449709). Digested muscles were harvested on culture dishes coated with 10μg mL^−1^ laminin (Sigma-Aldrich, Ref. 114956-81-9) and cultured with F-12 media (Ham’s F-12 Nutrient Mix, Gibco; Cat# 21127-022) complemented with 20% FBS, 10% horse serum, 1% L-glutamine, 1% penicillin/streptomycin and 2.5 ng/mL bFGF (Recombinant Human FGF-basic. Peprotech; Cat# 100-18B). For growing human primary muscle progenitors, cells were cultured in high-glucose DMEM (Sigma-Aldrich; Cat# D5671) supplemented with 20% FBS, 10% horse serum, 1% L-glutamine and 1% penicillin/streptomycin. For growing mouse primary muscle progenitors, cells were cultured in high-glucose DMEM supplemented with 10% FBS, 1% L-glutamine and 1% penicillin/streptomycin. For differentiation, both human and mouse primary muscle progenitor cells were cultured in high-glucose DMEM supplemented with 2% horse serum, 1% L-glutamine and 1% penicillin/streptomycin.

### Transfections and immunostaining

Transfections of human and mouse primary muscle progenitor cells were performed as previously described [34]. Briefly, primary muscle progenitor cells were transfected with 100nM of miR-24-3p mimic (Syn-mmu-miR-24-3p, Qiagen, MSY0000219), 100nM of miR-24-3p inhibitor (Anti-mmu-miR-24-3p, Qiagen, MIN0000219), 100nM of scrambled control (AllStars Negative Control siRNA Print, Qiagen, Cat. number.: 1027280) or 60nM of siRNA against Prdx6 (ThermoFisher Scientific, human: Ref. s18428; mouse: Ref. s67375) using Lipofectamine 2000 transfection reagent (ThermoFisher, Ref. 11668019). Culture media was changed to differentiation media (high-glucose DMEM complemented with 2% HS, 1% P/S, 1% α-glutamine) 6 hours after transfection. No media was changed until collection or staining of the cells. Cells transfected with the empty vector (lipofectamine control) and mock-transfected cells (scrambled control) served as controls. Immunostaining was performed 48 hr (Ki67 staining), 4 days (viability assay), 7 days (SA-β-galactosidase staining) and 7-10 days (MF 20 staining) after transfection. RNA and protein were isolated 48 hr after transfection. Staining for MF 20, SA-β-galactosidase, MF 20, Ki67 and viability assay was performed as previously published methods [34].

### miR:target binding reporter assay

5ʹUTR of Prdx6-202 transcript regions with either the wild-type or mutated miR-24-3p target sites were synthesized using GeneArt service (Thermo Scientific). The wild type or mutated sequences were subcloned into a GFP TOPO vector (Thermo Scientific). C2C12 myoblasts were cultured in 96-well plates and transfected using Lipofectamine 2000™ (Thermo Scientific) with either 200ng of the wild type or mutant sensor and with either 100nM of the miR scrambled control or 100nM miR-24 mimic. Each experiment was carried out using at least two independent plasmid preparations in triplicate. GFP fluorescence was measured 48 hr following transfections using FLUOstar Optima microplate reader (BMG Labtech).

### *In vitro* model of adaptive response to acute oxidative stress

Human and mouse primary muscle progenitor cells were cultured with growth media in 96-well plates. When 70-80% confluency was achieved, cells were transfected with 100nM microRNA mimic, anti-microRNA, scrambled control or 60nM siRNA against Prdx6 diluted in differentiation media as described previously, including 5-6 technical replicates per biological replicate. Media was removed 5 hours after transfection and cells were treated with 100μM of H_2_O_2_ dissolved in DPBS and then incubated in for 24 hours in a humidified 37°C, 5% CO_2_ incubator. After removing the H_2_O_2_ solution, cells were loaded with 10μM CM-H_2_DCFDA dissolved in DPBS and incubated in the dark for 30 minutes in a humidified 37°C, 5% CO_2_ incubator. Cells were then washed twice with DPBS and fluorescence intensity was measured using a FLUOstar OPTIMA microplate reader (BMG Labtech) for the detection of ROS (Emission: 517-527nm; excitation: 492-495nm). n=3 biological replicates per group with 5-6 independent technical replicates per individual.

### Image analysis

Cells were semi-automatically quantified using Fiji and ImageJ [35] followed by manual correction. At least 5 or more random images from different field of view per biological sample at 10x magnification (100x total magnification) were captured. A median total cell count of 1988 per condition was quantified in an independent way by three individuals. For myogenic differentiation analyses, fusion index is shown as the percentage of nuclei contained within myotubes to the total number of nuclei in each microscopic field. For the quantification of senescent cells, β-galactosidase activity values (BGAVs) were calculated as previously described by Shlush et al. [36]. Cells with a BGAV ≥ 15 were considered as highly senescent (SA-βgal^high^, characterised by an intense blue staining); cells with a BGAV between 5-14 both inclusive were considered as low senescent (SA-βgal^low^, characterised by an light blue staining); and cells with a BGAV < 5 were considered as non-senescent (SA-βgal^non^, no blue staining). All the immunostaining quantifications were manually curated. Images were captured using Nikon Eclipse Ti-E inverted confocal microscope (for MF 20, viability assay, Ki67 and CM-H_2_DCFDA) and Carl Zeiss Axiovert 200 inverted microscope (for SA-βgal staining).

### Statistical analysis

Details of the statistical analyses used per experiment are described in the corresponding figure legend. Two-tailed unpaired Student t-test, Mann-Whitney test or F-test were performed for the analysis of statistical differences between two groups as stated. One way or two-way A.N.O.V.A. followed by Tukey’s multiple comparison test, or Kruskal-Wallis followed by Dunn’s multiple comparison test with 95% Confidence Interval was performed to compare more than two groups as indicated. p-value < 0.05 was considered statistically significant. For the transfection experiments, individual values representing the same biological replicate have been matched with a dotted line for a better interpretation of the results. When appropriate, results were normalised to their respective control groups to acquire a more representative evaluation of the effects of the treatment. Statistical analysis were performed using GraphPad Prism version 8.4.2 for Windows (GraphPad Software, La Jolla California USA, www.graphpad.com).

### Gene ontology

A list of human and mouse miR-24 predicted targets were obtained from TargetScanHuman 6.2 [37–39]. Human and mouse miR-24:targets network interaction and GO analyses were performed using Cytoscape v.3.8.0 [40] and ClueGO v.2.5.6 plugin for Cytoscape [41], respectively. Details of the statistics used for ClueGO are specified in the corresponding figure legend. Generally: Enrichment/Depletion (Two-sided hypergeometric test); Minimum p-value cut-off=0.01; Correction Method=Bonferroni step down; Min GO Level=5; Max GO Level=8; Kappa Score Threshold=0.4-0.55.

## Results

### miR-24 is downregulated during skeletal muscle regeneration and ageing

The number of satellite cells and their function, which depends on their viability, myogenic potential and senescence, have been previously reported to be affected by ageing [9, 42–45]; however, the regulation of these processes is not well understood. We hypothesised that miRs regulate the decline in satellite cell function during ageing through controlling redox balance, as changes in redox signalling are known to control muscle regeneration [4]. Using targeted searches of miRs potentially regulating satellite cell ageing and redox homeostasis, we selected miR-24, previously shown to be regulated during satellite cell activation and ageing [20, 46], for further investigation. Some of the miR-24 putative targets in humans, analysed using TargetScan and ClueGO plugin for Cytoscape, were genes associated with the cellular response to oxidative stress **(Figure 1A)**. Among these targets was Prdx6, known to be involved in skeletal muscle adaptation under oxidative stress [47]. Multiple targets of miR-24 were also conserved between human and mouse **(Figure 6C)**. We therefore investigated changes in miR-24 expression in FACS-sorted satellite cells. Consistent with published data [9, 42–45], we observed a decrease in the total number of FACS-sorted satellite cells during ageing **(Figures 1B, S1)** and myogenic progenitors from old mice displayed increased senescence **(Figures 1C, 3A)**, reduced viability **(Figures 1D, 2A)**, reduced myogenic potential **(Figures 1E, 2D)** and increased ROS **(Figures 1F, 1G)**. miR-24 expression was downregulated in satellite cells from old mice **(Figure 1H)**. The expression of miR-24 was also analysed by RT-qPCR in an *in vivo* model of skeletal muscle regeneration following barium chloride (BaCl_2_) injection in the tibialis anterior (TA) **(Figures 1I, 1J)**. miR-24 basal levels were not altered in TA muscle during ageing, as opposed to its downregulation in satellite cells **(Figure 1H, 1J)**, but its expression was increased one day after muscle injury and returned to basal levels after seven days in the injured muscle of adult mice, suggesting a potential role of miR-24 in the early stages of muscle regeneration after acute injury. However, the expression of miR-24 did not increase in old mice following injury: miR-24 expression was significantly lower at 1-21 days after injury compared to the adult mice **(Figure 1J)**. These data suggest that the downregulation of miR-24 in satellite cells **(Figure 1H)** may be related to the age-related decline in satellite function and consequently muscle regeneration following acute injury.

**Figure 1.**
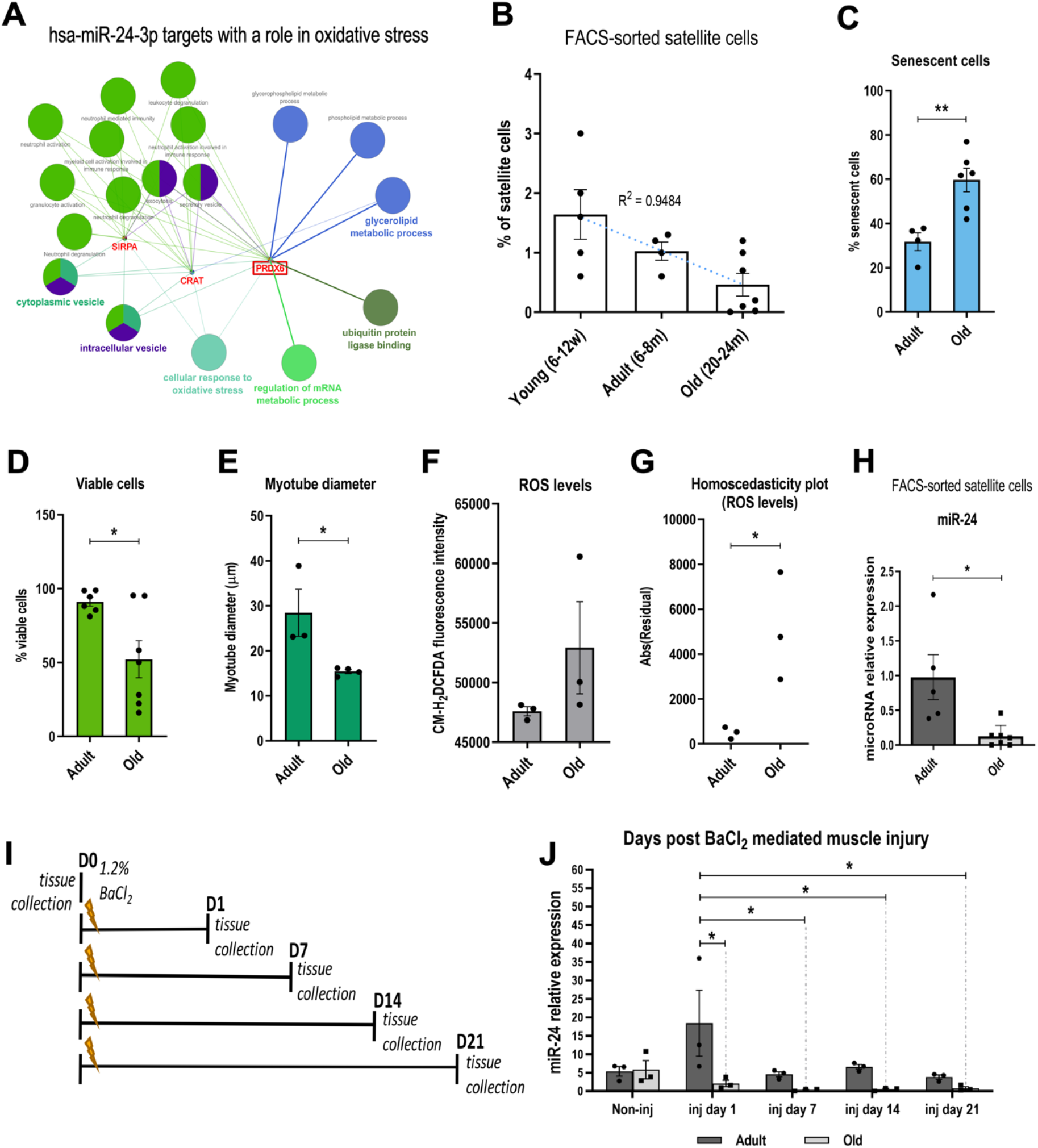
miR-24 expression is affected by muscle injury and ageing. **(A)** miR-24 is predicted to target genes and processes associated with redox balance in humans. Gene ontology (GO) analysis was performed ClueGO plugin for Cytoscape (v.2.5.6). Statistical test used for ClueGO: Enrichment/Depletion (Two-sided hypergeometric test). p-value cut-off=1.0E-4. Correction Method = Bonferroni step down. Min GO Level=5; Max GO Level=8; Number of Genes=16; Min Percentage=4.0; Kappa Score Threshold=0.4. Only targets involved in ‘cellular response to oxidative stress’ are shown. **(B)** The percentage of mouse satellite cells decreases during ageing (n=4-7, R^2^=0.9484). **(C-E)** Myogenic progenitors from old mice are less viable, have decreased myogenic potential and display increased senescence (n=3-6, two-tailed unpaired Student’s t-test). **(F)** The accumulation of ROS assessed using the CM-H_2_DCFDA assay. The graph shows mean fluorescence intensity values of CM-H_2_DCFDA in myogenic progenitors from adult and old mice. **(G)** Homoscedasticity plot showing increased levels of ROS in mouse myogenic progenitors during ageing (n=3, F-test for the comparison of unequal variances between the two groups). **(H)** qPCR showing decreased expression of miR-24 in mouse satellite cells during ageing. Expression relative to Snord61 is shown (n=5-7, Mann-Whitney test). **(I)** Diagram representing tissue collection points following mouse tibialis anterior (TA) muscle injury using BaCl_2_. **(J)** qPCR of miR-24 in the tibialis anterior muscle after injury with BaCl_2_ in adult and old mice. Expression relative to Snord61 is shown (n=3, two-way A.N.O.V.A followed by Tukey’s multiple comparison test with 95% Confidence Interval). For all the figures unless stated otherwise: Young: 6-12 weeks old; adult: 6-8 months old; old: 20-24 months old. For miR-24 qPCR in satellite cells: adult: 1-8 months old; old: 20-24 months old. p-value < 0.05 was considered as statistically significant (*, ^#^ p < .05; **, ^# #^ p < .01; ***, ^# # #^ p < .001). Error bars show S.E.M.

**Figure 2.**
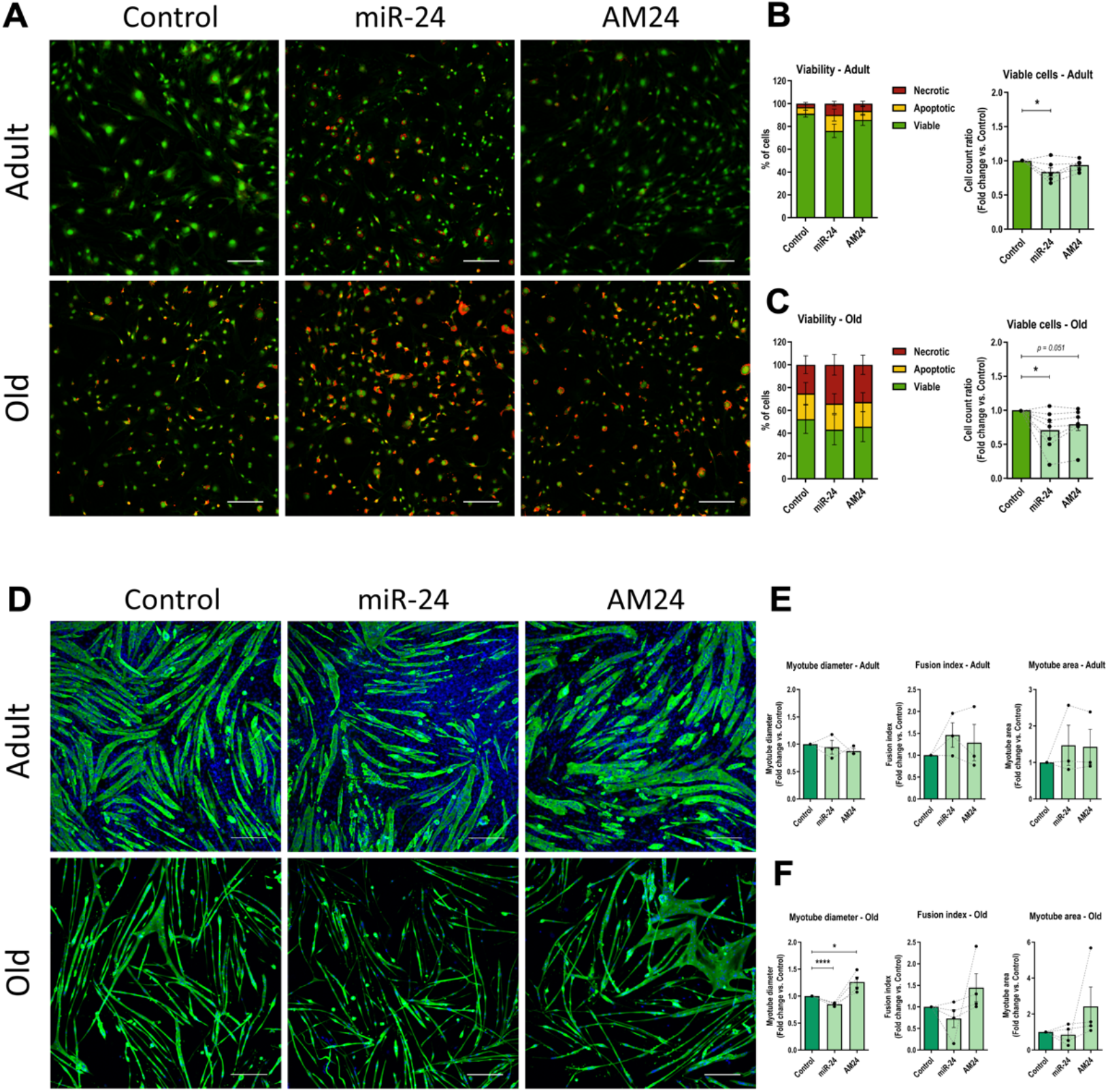
miR-24 regulates viability and differentiation of myogenic progenitors during ageing. **(A, D)** Myogenic progenitors isolated from adult and old mice were transfected with miR-24 or AM24. Cells transfected with the empty vector or scrambled control were used as control (Control). Scale bars: 200μm. **(A)** Viability assay was performed with ethidium bromide and acridine orange for the assessment of viable (green), apoptotic (yellow) and necrotic (red) cells. **(B, C)** miR-24 overexpression resulted in decreased viable cells from both adult and old mice. miR-24 inhibition also affected viability of cells isolated from old mice (n=6-7, two-tailed unpaired Student’s t-test compared to control). **(D)** MF 20 (anti-myosin heavy chain; green) and DAPI (blue) immunostaining were performed for myogenic differentiation and nuclei identification, respectively. **(E)** Overexpression or inhibition of miR-24 did not significantly affect the differentiation of myogenic progenitors from adult mice. **(F)** miR-24 overexpression resulted in decreased myotube diameter whereas miR-24 inhibition resulted in bigger myotubes in myogenic progenitors from old mice compared to Control group (n=3-4, two-tailed unpaired Student’s t-test compared to control). For all the figures: adult: 6-8 months old; old: 20-24 months old. p-value < 0.05 was considered as statistically significant (*p < .05; **p < .01; ***p < .001; ****p < .0001). Error bars show S.E.M.

**Figure 3.**
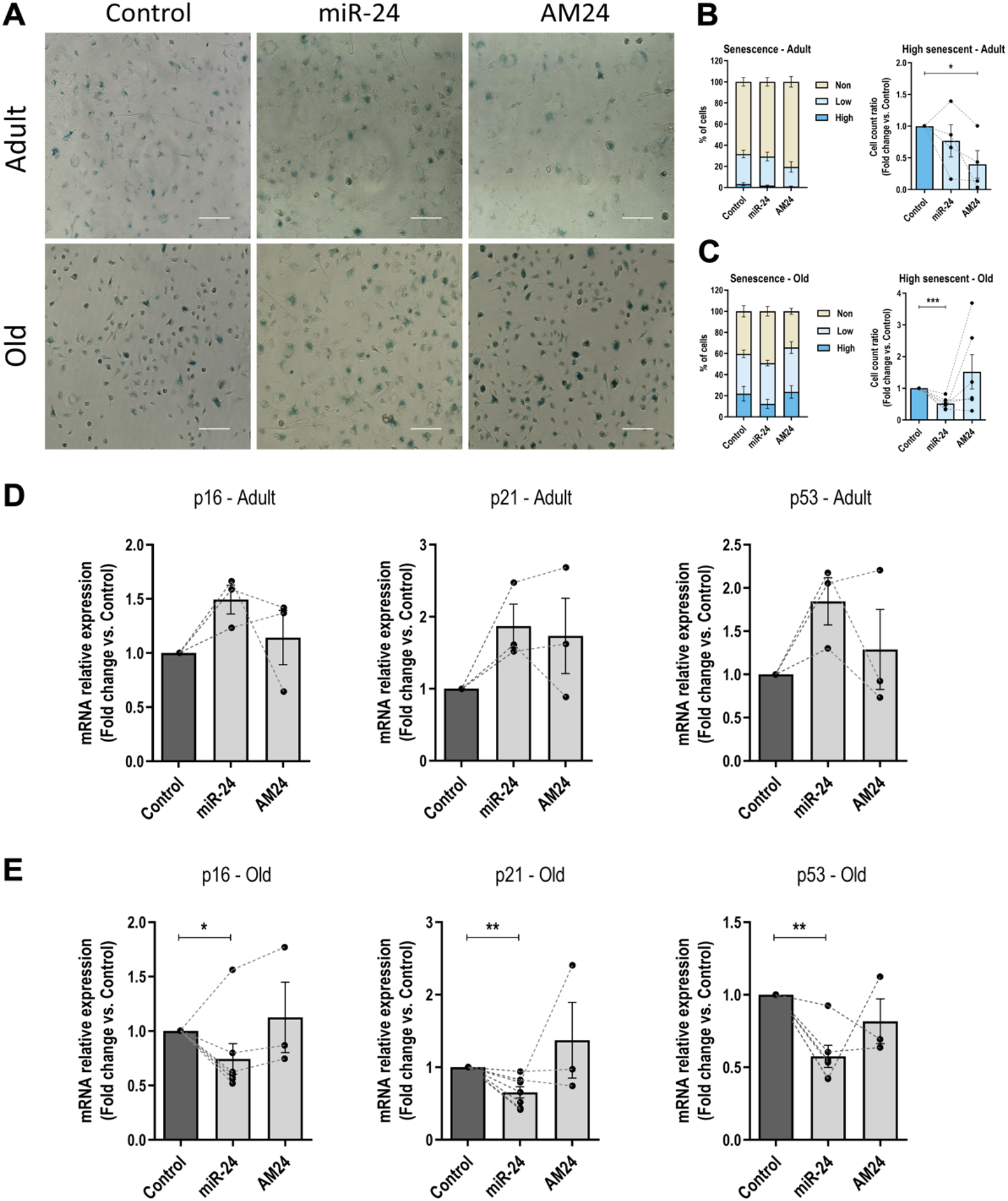
miR-24 regulation of senescence of myogenic progenitors is age-dependent. **(A)** Myogenic progenitors isolated from adult and old mice were transfected with miR-24 mimic (miR-24) or anti-miR (AM24). Cells transfected with the empty vector or scrambled control were used as control group (Control). SA-βgal staining was performed for the assessment of senescent cells (blue). Scale bars: 200μm. **(B, C)** miR-24 inhibition resulted in fewer highly senescent cells in the adult **(B)**, whereas miR-24 overexpression resulted in fewer highly senescent cells in the old mice **(C)** (n=4-6, two-tailed unpaired Student’s t-test compared to control). **(D, E)** The expression of senescence-associated genes was not significantly changed by miR-24 overexpression in myogenic progenitors from adult mice **(D)**, whereas p16, p21 and p53 expression was significantly downregulated following miR-24 overexpression in myogenic progenitors from old mice **(E)** (n=3-7, Kruskal-Wallis test followed by Dunn’s multiple comparisons test with 95% Confidence Interval; expression relative to 18S is shown). For all the figures: adult: 6-8 months old; old: 20-24 months old. p-value < 0.05 was considered as statistically significant (*p < .05; **p < .01; ***p < .001). Error bars show S.E.M.

### miR-24 regulates viability and myogenic potential of myogenic progenitors during ageing

The function of satellite cells in muscle regeneration depends on their viability and myogenic potential. Our data demonstrated that both are affected by ageing **(Figures 1D, 1E)**. To determine the physiological consequences of miR-24 dysregulation in satellite cells during ageing and regeneration, myogenic progenitors isolated from adult and old mice were transfected with miR-24 mimic, AM24 or mock-transfected (Control) **(Figures 2A, 2D)**. Myoblasts were stained to evaluate differentiation (MF 20), proliferation (Ki67) and viability **(Figures 2, S4)**. miR-24 had no significant effects on myogenic progenitor proliferation **(Figure S4)**. However, overexpression of miR-24 resulted in the presence of more necrotic and apoptotic myogenic progenitors from both adult and old mice **(Figures 2B, 2C)**.

miR-24 overexpression or inhibition in myogenic progenitors from adult mice had no effects on their myogenic differentiation **(Figure 2E)**. Inhibition of miR-24 in myogenic progenitors from old mice had a higher fusion index and increased myotube area, although this was not statistically significant **(Figure 2F)**. However, miR-24 overexpression in myogenic progenitors from old mice resulted in the presence of smaller myotubes, whereas inhibition of miR-24 resulted in the presence of larger myotubes only in cells isolated from old mice **(Figure 2F)**.

### miR-24 may differentially regulate senescence of myogenic progenitors from adult and old mice

Satellite cells have been previously shown to undergo senescence during ageing, suggested to affect their function [48, 49]. Moreover, miR-24 has been previously demonstrated to regulate the expression of p16 in human diploid fibroblasts and cervical carcinoma cells [28, 50]. In order to determine whether miR-24 regulates cellular senescence, myogenic progenitors from adult and old mice were transfected with miR-24 mimic or anti-miR (AM24), and senescence phenotype was examined by senescence-associated β-galactosidase staining (SA-βgal) and expression of tumour suppressor genes **(Figure 3)**. In myogenic progenitors from adult mice, overexpression of miR-24 had no effect on the number of highly senescent cells (SA-βgal^high^), characterised by an intense SA-βgal blue staining) or overall proportions of senescent cells as compared to control, whereas miR-24 inhibition led to a decreased number of SA-βgal^high^ cells as compared to control **(Figure 3B)**. Moreover, the expression of genes associated with cell senescence p16, p21 and p53 were not significantly changed in myogenic progenitors from adult mice following miR-24 overexpression or inhibition **(Figure 3D)**. In myogenic progenitors from old mice, which express lower levels of miR-24 compared to adult mice, miR-24 overexpression led to a decrease in the number of SA-βgal^high^ cells as compared to control cells **(Figure 3C)**. This was consistent with the downregulation of p16, p21 and p53 expression following miR-24 overexpression **(Figure 3E)**. However, inhibition of miR-24 in myogenic progenitors from old mice had no significant effect on number of SA-βgal^high^ cells or the overall proportion of senescent cells **(Figure 3C)**; this was consistent with no significant changes in senescence-associated genes **(Figure 3E)**. This could be associated with already low levels of miR-24 in cells from older mice. The differences in phenotypic changes following miR-24 overexpression or inhibition on myogenic progenitor senescence from adult and old mice are consistent with those in myogenic progenitor viability and myogenic potential, which suggests a context-dependent function of miR-24.

### miR-24 directly regulates the expression of Prdx6 in myogenic progenitors

To determine whether miR-24 may regulate muscle regeneration through regulation of viability, differentiation and senescence of myogenic progenitors from old mice, we focused on redox-associated target genes as redox homeostasis has been shown to regulate all these processes during ageing [4]. Prdx6 has been shown to regulate skeletal muscle adaptation under increased oxidative stress [47] and miR-24 was previously shown to target Prdx6 in humans **(Figure 1A)** [51]. We therefore hypothesised that miR-24 may regulate the viability and myogenic potential of satellite cells by controlling the expression of Prdx6 during ageing. Prdx6 was upregulated in FACS-sorted satellite cells isolated from old mice compared to adult mice **(Figure 4A)**, but was not significantly upregulated in the tibialis anterior muscle from old mice compared to adult mice **(Figure S3);**this is consistent with downregulation of miR-24 expression in satellite cells but not muscle during ageing **(Figure 1H, J)**. To determine whether upregulation of Prdx6 in satellite cells may be associated with miR-24 downregulation, we analysed the sequence of mouse Prdx6 for miR-24 binding sites. A target site for miR-24 was found between position 2-6 (6mer) of the mature microRNA-24 and position 41-45 5’UTR of the mouse Prdx6-202 transcript **(Figure 4B and materials and methods)**. A GFP reporter containing the wild type or mutated miR-24 binding site for miR-24 was generated **(Figure 4C)**. C2C12 myoblasts were transfected with reporter constructs containing wild type or mutated miR-24 binding site within the Prdx6 5’UTR fragment and co-transfected with either miR-24 mimic or scrambled sequence used as a control. GFP levels were decreased in the cells transfected with wild type construct co-transfected with miR-24 as compared to scrambled control (Scr) treated cells, but not in the cells treated with the mutated construct co-transfected with miR-24 or control Scr microRNA, indicating that miR-24 directly binds to Prdx6 mRNA in mouse myoblasts **(Figure 4C)**. We next investigated Prdx6 expression following miR-24 overexpression or downregulation in myogenic progenitors from adult and old mice. miR-24 did not significantly change the levels of Prdx6 mRNA in myogenic progenitors from adult mice **(Figure 4D)**. However, Prdx6 expression was significantly downregulated following miR-24 overexpression in myogenic progenitors from old mice **(Figure 4E)**. These data suggest that the regulation of Prdx6 by miR-24 may depend on the cellular context, for example different levels of expression of its target genes associated with ageing. In this case, as Prdx6 levels are increased during ageing, Prdx6 may be more susceptible to regulation by miR-24 in myogenic progenitors from old mice. Alternatively, the presence or absence of other factors regulating Prdx6 expression upstream of miR-24 during ageing, may contribute to the level of Prdx6 expression regulation by miR-24.

**Figure 4.**
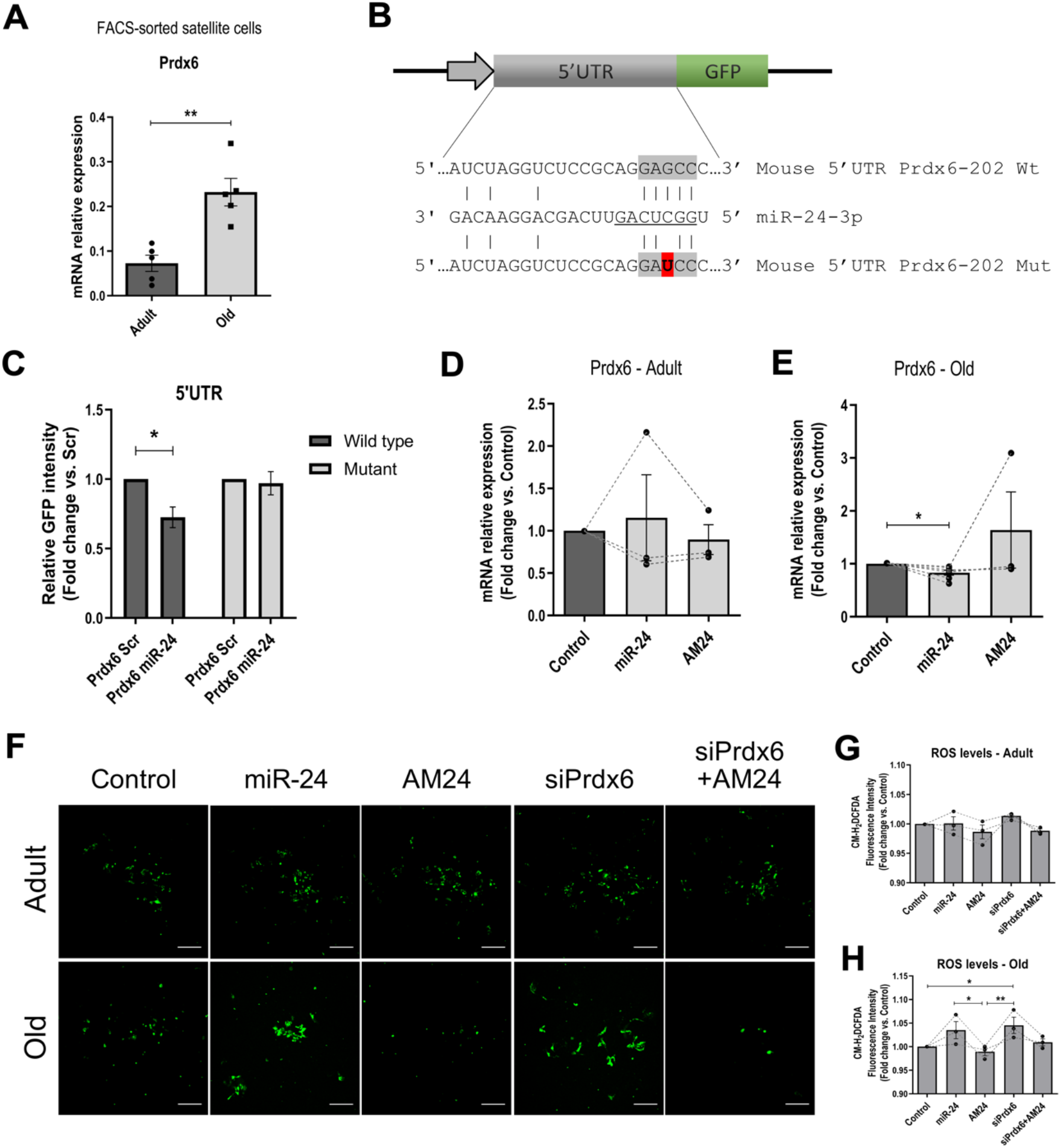
miR-24:Prdx6 interactions regulate ROS production in an *in vitro* model of acute oxidative stress. **(A)** qPCR showing Prdx6 expression in mouse satellite cells during ageing. Expression relative to beta-actin is shown. Adult: 1-8 months old. Old: 20-24 months old (n=5, Mann-Whitney test). **(B)** Putative miR-24-3p seed sequence in the 5’ UTR of Prdx6 gene (highlighted in grey). Mutated seed sequence used for 5’UTR microRNA:mRNA target interaction is shown. Mutation is shown in red. **(C)** miR-24 directly regulates the expression of Prdx6. GFP-Prdx6 5’UTR sensor construct containing the wild type or mutated seed sequence were transfected into C2C12 myoblast cell line and co-transfected with miR-24 or scrambled control (Scr). The wild type construct transfected with miR-24 mimic shows less GFP fluorescence intensity compared to the scrambled control, but not in the mutated construct. (Representative data shown; n=3, two-tailed unpaired Student’s t-test). **(D, E)** qPCR showing the expression of Prdx6 after microRNA mimic or antagomiR (AM24) transfection in primary myogenic progenitors isolated from adult **(D)** and old mice **(E)**. Expression relative to 18S is shown. Adult: 6-8 months old; old: 20-24 months old (n=3-7, Kruskal-Wallis test followed by Dunn’s multiple comparisons test with 95% Confidence Interval). **(F)** CM-H_2_DCFDA staining for the assessment of ROS generation in myogenic progenitor cells. Myogenic progenitors isolated from adult and old mice were transfected with miR-24 mimic (miR-24), anti-miR (AM24) and/or siRNA against Prdx6 (siPrdx6) as indicated. Cells transfected with scrambled control were used as control group (Control). After 5 hours, media was changed and cells were treated with 100μM H_2_O_2_ for 24 hours to mimic oxidative stress. Followed H_2_O_2_ treatment, CM-H_2_DCFDA staining was performed as an indicator for reactive oxygen species (ROS) generation. Scale bars: 200μm. Adult: 6-8 months old. Old: 20-24 months old. **(G, H)** ROS fluorescence intensity quantification using the mean of 5-6 repeated independent technical measures from 3 biological replicates (n=3, repeated measures one-way A.N.O.V.A. followed by Tukey’s multiple comparison test with 95% Confidence Interval). For all the figures unless stated otherwise: adult: 6-8 months old; old: 20-24 months old. p-value < 0.05 was considered as statistically significant (*p < .05; **p < .01; ***p < .001). Error bars show S.E.M.

### Inhibition of miR-24 reduces ROS production during ageing through Prdx6 de-repression in an *in vitro* model of oxidative stress

As PRDX6 is a well-known antioxidant protein, downregulation of miR-24 and consecutive upregulation of Prdx6 in satellite cells from old mice may serve as a protective mechanism against an increased oxidative environment after acute injury.

To test this hypothesis, we used an *in vitro* model to mimic acute oxidative stress. Primary myogenic progenitors isolated from adult and old mice were transfected with miR-24 mimic, anti-miR (AM24), siRNA against Prdx6 (siPrdx6), co-transfected with both anti-miR and siRNA against Prdx6 (AM24+siPrdx6). After transfection, media was removed and cells were incubated with 100μM H_2_O_2_ for 24 hours. ROS generation was measured the following day using CM-H_2_DCFDA fluorescence staining **(Figure 4F)**. ROS generation in myogenic progenitors from adult mice was not changed following miR-24 overexpression or inhibition, or Prdx6 downregulation **(Figure 4G)**. However, ROS generation in myogenic progenitors from old mice was increased following Prdx6 silencing as compared to control cells (Scr) **(Figure 4H)**. In addition, lower ROS levels were detected following miR-24 inhibition in comparison to miR-24 overexpression and Prdx6 silencing **(Figure 4H)**. In order to determine whether the effect of miR-24 inhibition on ROS generation could be mediated by changes in PRDX6 levels, myogenic progenitors were co-transfected with AM24 and siPrdx6. Co-transfection of AM24 together with siRNA against Prdx6 counteracted the increased ROS generated by lower Prdx6 levels in myogenic progenitors from old mice **(Figures 4F, 4H)**. Together, these data suggest that miR-24 may regulate Prdx6 anti-oxidant activity via regulating its expression in myogenic progenitors from old mice. These data are consistent with high levels of miR-24 decreasing myotube diameter and cell viability during ageing **(Figures 2A, 2D)**.

### miR-24 regulation of myogenic potential and viability by controlling Prdx6 is conserved inhuman primary myogenic progenitors

We further explored whether miR-24 has the same phenotypic effects in human myogenic progenitors as in murine myogenic progenitors. Primary myogenic progenitor cells isolated from young adults were transfected with miR-24 mimic, AM24 or scrambled or empty vector control. Differentiation, viability, senescence and proliferation were analysed as previously for murine cells **(Figure 5A)**. miR-24 overexpression resulted in decreased myotube diameter and myogenic area **(Figure 5B)**, and decreased number of viable cells **(Figure 5C)**, consistently with the effects of miR-24 in murine cells. Interestingly, miR-24 overexpression led to an increased number of senescent cells **(Figure 5D)**as compared to control group, again highlighting that miR-24 regulation of senescence or the lack of it may depend on the context, such as age or species. Ki-67 immunostaining revealed no significant effect on proliferation after overexpression or inhibition of miR-24 in human primary myogenic progenitors **(Figure S4)**, consistently with the lack of miR-24-mediated regulation during proliferation in murine myogenic progenitors.

**Figure 5.**
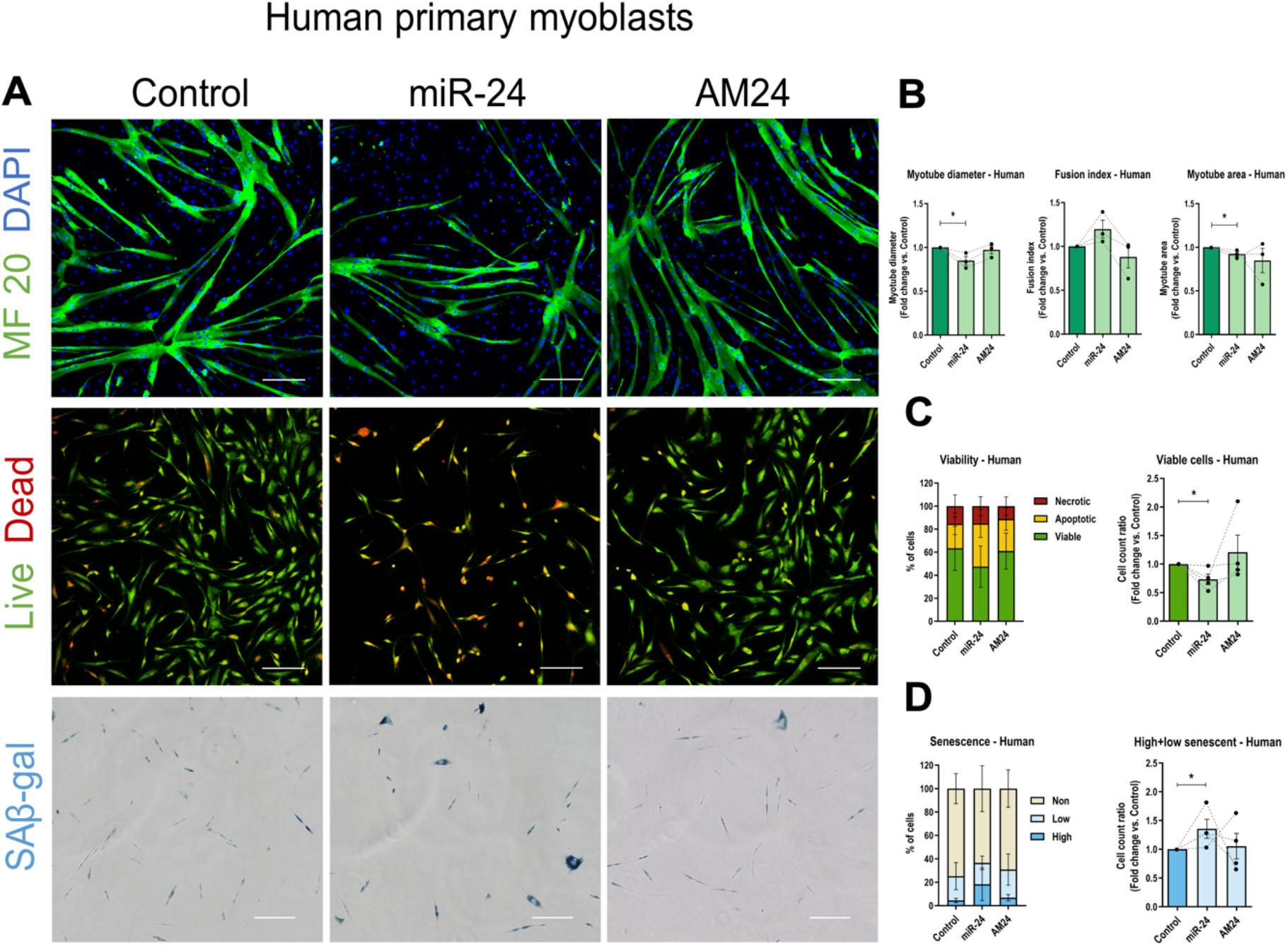
miR-24 regulates differentiation, viability and senescence of human primary myogenic progenitors. **(A)** Human primary myogenic progenitor cells isolated from young-adult individuals were transfected with miR-24 or AM24. Cells transfected with the empty vector or scrambled control were used as control (Control). MF 20 (anti-myosin heavy chain; green) and DAPI (blue) immunostaining were performed for myogenic differentiation and nuclei identification, respectively. Viability assay was performed with ethidium bromide and acridine orange for the assessment of viable (green), apoptotic (yellow) and necrotic (red) cells. Senescence Associated β-galactosidase (SA-βgal) staining was performed for the assessment of senescent cells (blue). Scale bars: 200μm. **(B-D)** miR-24 overexpression resulted in thinner myotubes, less viability and increased number of senescent cells (n=3-4, two-tailed unpaired Student’s t-test compared to Control group). p-value < 0.05 was considered as statistically significant (*p < .05; **p < .01; ***p < .001). Error bars show S.E.M.

As Prdx6 was previously shown to be a target of miR-24 in human cells [51], we determined whether miR-24:Prdx6 interactions were involved in regulating ROS levels in human cells. Primary myogenic progenitors isolated from young adults were transfected with either miR-24 mimic, anti-miR (AM24), siRNA against Prdx6 (siPrdx6), co-transfected with both anti-miR and siRNA against Prdx6 (AM24+siPrdx6) or transfected with the scrambled control (Scr). Media was changed 5 hours after transfection and cells were treated with 100μM H_2_O_2_ diluted in DPBS for 24 hours. Subsequently, ROS generation was measured using CM-H_2_DCFDA fluorescence staining **(Figure 6A)**. Similar to the results obtained from myogenic progenitors isolated from old mice **(Figures 4F, 4H),** higher ROS levels were detected after miR-24 overexpression and Prdx6 silencing in comparison to miR-24 inhibition compared to scrambled control group. ROS levels were also significantly reduced after miR-24 inhibition compared to control group. In addition, co-transfection of AM24 together with siRNA against Prdx6 also counteracted the increased ROS generated by lower Prdx6 levels **(Figure 6B).**

**Figure 6.**
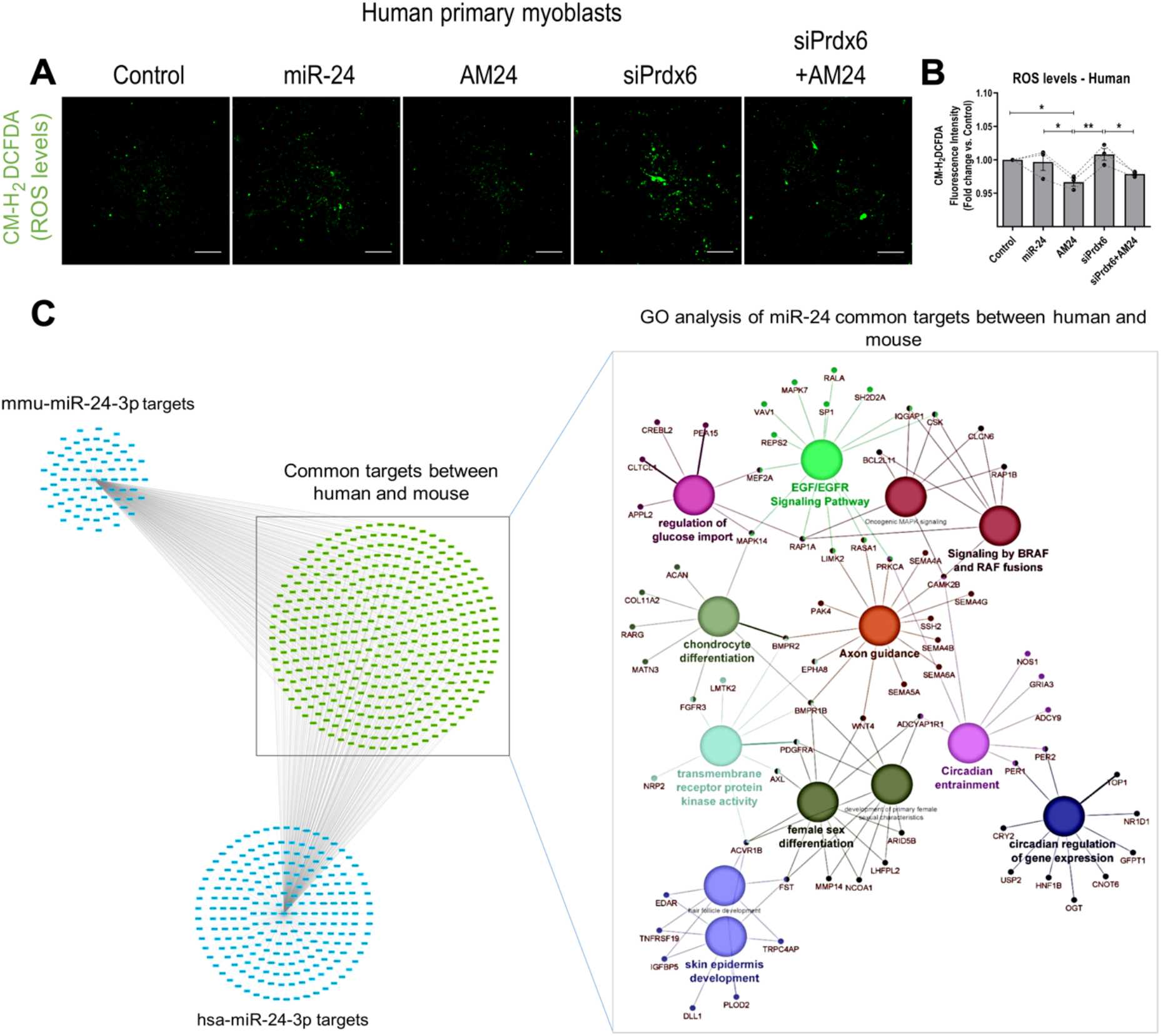
miR-24:Prdx6 interactions regulate ROS production in human primary myogenic progenitors. **(A)** Human primary myogenic progenitor cells were cultured and transfected with miR-24 mimic (miR-24), anti-miR (AM24) and/or siRNA against Prdx6 (siPrdx6) as indicated. Cells transfected with scrambled control were used as control group (Control). Media was removed 5 hours after transfection and cells were treated with 100μM H_2_O_2_ for 24 hours to mimic oxidative stress. After H_2_O_2_ treatment, CM-H_2_DCFDA staining was performed as an indicator for reactive oxygen species (ROS) production in the cells. Scale bars: 200μm. Human: young-adult. **(B)** ROS fluorescence intensity quantification using the mean of 5-6 repeated independent technical measures from 3 biological replicates (n=3, repeated measures one-way A.N.O.V.A. followed by Tukey’s multiple comparison test with 95% Confidence Interval). p-value < 0.05 was considered as statistically significant (*p < .05; **p < .01; ***p < .001). Error bars show S.E.M. **(C)** Gene ontology analysis revealing the main biological processes in miR-24 common predicted targets between humans and mice. Network and GO analysis were performed using Cytoscape (v.3.8.0) and ClueGO plugin for Cytoscape (v.2.5.6), respectively. Statistical Test used for ClueGO: Enrichment/Depletion (Two-sided hypergeometric test). p-value cut-off=0.01. Correction Method=Bonferroni step down. Min GO Level=5; Max GO Level=8; Number of Genes=7; Min Percentage=8.0; Kappa Score Threshold=0.55.

These data show the consistent regulation of myogenic potential and viability of myogenic progenitors from humans and old but not adult mice, and consistent but stronger miR-24:Prdx6-mediated regulation of ROS production in human as compared to mouse myogenic progenitors. This may be due to higher levels of Prdx6 in myogenic progenitors from old mice as compared to adult mice and/or a stronger miR-24 interaction to the 3’UTR Prdx6 binding site in humans as compared to the 5’UTR in mice.

## Discussion

Muscle ageing comprises the disruption of a wide range of physiological processes, including the loss of muscle mass and strength, fatty degeneration, impaired glucose metabolism, altered ROS balance, loss of neuromuscular junctions and stem cell exhaustion and senescence. These processes ultimately affect the myocyte niche compromising satellite cell functionality and regenerative potential in response to injury [52].

Following injury and during the regenerative process, an acute increase in endogenous ROS is required to promote a pro-inflammatory environment that helps with macrophage recruitment, allowing an array of adaptive responses and signalling events such as during exercise-induced ROS generation, wound repair and inflammation [53–55]. Following an acute increase, ROS levels decrease at later stages of regeneration to allow muscle hypertrophy and remodelling during the anti-inflammatory phase [55–57]. However, ROS levels must be tightly regulated, as chronically elevated ROS may induce irreversible protein modifications, aberrant signalling, DNA damage and mutagenesis [58]. When damage persists, cellular stressors can trigger a transient cell cycle arrest via activation of p53/p21 or p16/pRB axes, which might eventually result in the induction of cellular senescence or cell death programs such as apoptosis and autophagy [59].

This study aimed to investigate the underlying biological mechanisms of the microRNA miR-24-3p in muscle regeneration during ageing. Our results demonstrate an increased expression of miR-24 one day after acute injury in an *in vivo* model of skeletal muscle regeneration in adult mice. miR-24 expression returned to baseline levels 7 days after injury, when myoblasts stop proliferating and start differentiating to initiate tissue remodelling (from 7 to 21 days after injury in mice) [60] **(Figure 1J)**. We have also identified an upregulation of the anti-oxidant Prdx6 in mouse quiescent satellite cells during ageing **(Figures 4A),** and confirmed Prdx6 as a direct target gene of miR-24 in mice **(Figure 4C).** Using an *in vitro* model to mimic oxidative stress, we have demonstrated that inhibition of miR-24 counteracts ROS through Prdx6 de-repression in primary myogenic progenitors isolated from old mice **(Figures 4F, 4H)** . Our results suggest that downregulation of miR-24 in satellite cells and subsequent upregulation of its target gene Prdx6, has a protective role against a disrupted redox environment during ageing through preserving cell viability and their myogenic potential, but consequently disrupts the adaptive redox signalling required during muscle regeneration. Since miR-24 regulates the levels of senescence-associated p16, p21 and p53 (**Figure 3E**), downregulation of miR-24 may ultimately result in increased senescence of satellite cells resident in the muscle of old mice.

It has previously been reported that miR-24 is upregulated by HIF-1α (hypoxia inducible factor 1 alpha) in cardiomyocytes, protecting these cells from ischaemic injury [61]. In addition, miR-24 is known to inhibit the degradation of HIF-1α by targeting FIH1 (factor inhibiting HIF-1) resulting in a positive feedback loop [62]. It is possible that injury promotes the upregulation of both HIF-1α and miR-24 to induce early stages of muscle regeneration. However, miR-24 is not upregulated after injury in muscle from old mice **(Figure 1J)**. These findings suggest an impaired muscle adaptation in response to injury in the old mice, which might explain in part the defective regeneration associated with ageing. Moreover, miR-24 is downregulated only in satellite cells during ageing **(Figure 1H)**, but not in the whole tibialis anterior muscle under quiescent conditions **(Figure 1J)**. It is possible that aged satellite cells are more sensitive to ROS, and that an altered redox balance in the myofibre microenvironment during ageing can disrupt the expression of miR-24 in satellite cells. By using *in vitro* functional studies, miR-24 overexpression resulted in a higher number of necrotic and apoptotic myogenic progenitors with less viable cells in both adult and old mice **(Figures 2A, 2B, 2C)**. In addition, miR-24 inhibition resulted in bigger myotubes from old mice **(Figures 2D, 2F)**. It remains to be established whether the effect of miR-24 on myogenic differentiation is an indirect effect of its regulation of myogenic progenitor viability, and therefore more cells available to fuse, or whether miR-24 directly regulates myoblast fusion and maturation.

Interestingly, miR-24 had different effects on cellular senescence on myogenic progenitors from adult and old cells and human cells. It has been previously shown that senescent cells are generally characterised by a maintained expression of tumour suppressors such as p16^INK4a^, p21 and p53 [63]. The downregulation in the number of senescent cells and lower expression of senescence-associated factors in myogenic progenitors from old mice following miR-24 overexpression is consistent with miR-24 targeting p16, p21 and p53 **(Figures 3A, 3C, 3E)**[28, 29, 50]. Interestingly, primary myoblasts isolated from adult mice had fewer senescent cells after miR-24 inhibition **(Figures 3A, 3B)**. According to previous published data, miR-24 regulates the expression of tumour suppressor/senescence-associated proteins differently depending on the cell type and metabolic state of the cell: it reduced P16 protein levels but not p16^Ink4a^ mRNA levels in human diploid fibroblasts and cervical carcinoma cells [28]; and in chondrocytes, downregulation of miR-24 triggered p16 upregulation [29]. miR-24 also inhibited H2Ax in terminally differentiated hematopoietic cells making them vulnerable to DNA damage [30]. On the other hand, it has been also shown that miR-24 increased the expression of p53 while decreasing the levels of DNA topoisomerase 1 in fibroblasts isolated from newborn foreskin, resulting in oxidative stress-induced cellular senescence [31]. miR-24 also increased p53 and p21 protein levels in different cancer cell lines [64] and it induced p53 expression in human lens epithelial cells during ageing and oxidative stress [65]. Together, these results suggest that miR-24 exerts either an inhibitory or enhancer function over tumour suppressor/senescence-associated proteins depending on the cell cycle state, which is consistent with our findings. Context-dependent role of miRs has been previously demonstrated, as well as their dose-dependent regulation of physiological processes [66–69]. Moreover, miR-24 may regulate the expression of senescence-regulated genes via an upstream regulatory factor not yet identified by us. Of note, it has been demonstrated that miR-24 induces G2/S cell cycle arrest independently of p53 and p21 function through targeting DHFR in colorectal cancer cells, and despite the lower proliferative rate (mostly by cytotoxic effect) the surviving cells presented a more differentiated-like phenotype [64]. This might explain the higher trend in fusion index regardless of the lower cell viability after miR-24 overexpression in human primary myoblasts **(Figure 5A, 5B)**, as the same pathways involved in apoptosis are also known to regulate early myogenic differentiation. Moreover, Caspase-2 is transitionally increased at the onset of myogenesis, allowing cell cycle exit and myocyte differentiation through p21 activation [70] and adding apoptotic myoblasts to healthy myoblasts promoted the fusion of the viable cells [71]. The delicate balance between apoptotic, anti-apoptotic, proliferative and cell cycle arrest signals will ultimately determine whether some cells successfully differentiate/self-renew or, in contrast, die/become senescent. It is worth mentioning that cellular senescence analyses must be interpreted with caution. SA-β-gal activity derives from an increased activity of the lysosomal protein β-D-galactosidase in response to autophagy stimuli [72], and increased SA-β-gal and higher levels of tumour suppressor proteins may not always represent a permanent cell cycle arrest and cellular senescence *per se*, but can be sign of alternative processes involving a transient cellular arrest [73].

miR-24 has been previously reported to inhibit proliferation, migration and invasion while promoting apoptosis in human gastric cancer cells by targeting Prdx6 [51]. Similar to the results presented here in muscle regeneration after acute injury **(Figure 1J)**, miR-24 expression is dynamically changed during gastric metastasis progression [51]. Li *et al.* suggests that miR-24 is downregulated before metastasis to promote Prdx6 expression to cope with the excess ROS and prevent DNA damage. Moreover, the diverse activities of Prdx6, including peroxidase, PLA2 phospholipase and LPCAT activities, mean that it could potentially regulate different metabolic signalling pathways, from cell cycle, membrane repair and antioxidant response [74, 75]. For instance, the peroxidase activity of PRDX6 provides protection after H_2_O_2_ induced oxidative stress, but the PLA2 phospholipase activity is involved in NOX2 activation and subsequent generation of superoxide [75–77]. Previously, we have detected redox specific changes in the catalytic Cys47 of PRDX6 in peripheral nerves from old mice and the SOD1^−/−^ murine model of ageing [78, 79]. Whether these redox specific modifications result in a shift from less peroxidase activity to more PLA2 phospholipase activity of PRDX6 during ageing, which would lead to an increased pro-apoptotic role of PRDX6, remains to be determined [77]. Taking into account our data and the widely diverse results existing in the literature, we propose that miR-24, similar to its target Prdx6, displays a dual role depending on the cellular environment/context.

One limitation of this study is the variability in the efficiency of the transfection experiments. For the qPCR experiments, only samples where transfection efficiency was validated, either by an increased or inhibited expression of miR-24, were taken into consideration for the quantification of the target genes expression. In particular, the inhibition of miR-24 was challenging to achieve, which resulted in a reduced number of the biological replicates used for this particular group. Likewise, the efficiency in the transfection of the cells used for the immunofluorescence experiments might also be affected. Another limitation to be considered in this study is the use of solely one technique for the assessment of ROS generation and oxidative stress. Future studies should corroborate these findings using additional approaches. Similarly, SA-β-gal results must be interpreted with caution, as increased intensity of the staining in some cells did not always correlated with an extension of the cytoplasm, which is a well-known characteristic of senescent cells. Therefore, SA-β-gal staining and higher levels of tumor suppressor proteins may not always indicate a permanent cell cycle arrest, but probably a stress-induced transient cell cycle arrest that might trigger alternative processes such as apoptosis or autophagy. Noteworthy, myogenic progenitors isolated from mouse were from males whereas human samples were retrieved from female donors. Several studies have shown biological differences between rodent males and females in the development of sarcopenia and efficiency in muscle regeneration, as well as in the global expression of microRNAs in human skeletal muscle [80–83]. It is thus important to point out that the altered expression of genes in satellite cells and muscle progenitors shown in this study might be gender-specific in addition to species-associated differences.

In summary, our results identify a role for miR-24-3p through inhibition of Prdx6 in satellite cells during ageing which may play a key role in early stages of skeletal muscle regeneration after acute injury, through controlling adaptive redox and apoptotic signalling pathways. Moreover, our findings show that miR-24 regulation occurs in myogenic progenitors from humans and old, but not to the same extent in adult mice, and this is associated with miR-24 having more pronounced effects on ROS generation and myoblast differentiation during ageing. This mechanism may not be as strongly conserved in mice as in humans as miR-24 regulated ROS, myoblast viability, differentiation and senescence in myoblasts from adult humans. This is not surprising, as miR-24 binding site in Prdx6 resides at the 3’UTR of the Prdx6 transcript in humans, whereas in mice, this site has a weaker interaction at the 5’UTR of the Prdx6 transcript. The role of miR-24 in the regulation of cellular senescence requires further studies using a wider plethora of senescence markers and approaches.

In summary, we propose that miR-24:Prdx6 interactions might serve as an antioxidant protection mechanism against increased oxidative damage in muscle stem/progenitor cells during ageing, however, at the cost of their regenerative capacity. This mechanism is primarily mediated by conserved regulation of myogenic progenitor viability and their ability to differentiate and appears to be associated with regulation of cellular senescence during ageing.

---

**Figure S1.** Illustration showing the isolation of quiescent satellite cells by FACS (fluorescence activated cell sorting). The satellite cells population was positive for CD34, highly-positive for alpha7-integrin, and negative for Sca1, CD31 and CD45.

**Figure S2.** qPCR showing the expression of miR-24 after microRNA mimic or antagomiR (AM24) transfection in primary myogenic progenitors isolated from adult and old mice. Expression relative to Snord61 is shown (n=3-7, adult: 6-8 months old; old: 20-24 months old). F-test compared to control. p-value < 0.05 was considered as statistically significant (^#^ p < .05; ^# #^ p < .01; ^# # #^ p < .001; ^# # # #^ p < .0001). Error bars show S.E.M.

**Figure S3.** qPCR showing Prdx6 expression in mouse tibialis anterior muscle during ageing. Expression relative to ß-2 microglobulin is shown (n=3-4, adult: 6-8 months old; old: 20-24 months old). Mann-Whitney test (* < .05; **p < .01; ***p < .001). Error bars show S.E.M.

**Figure S4.** Ki67 staining of primary myogenic progenitors. Primary myoblasts isolated from adult and old mice and humans were transfected with miR-24 mimic (miR-24) for *in vitro* overexpression or antagomiR (AM24) for microRNA inhibition. Cells transfected with the empty vector or scrambled control were used as control group (Control) (**A**). Proliferation, shown as Ki67 positive cells (green), was not significantly affected by the overexpression or inhibition of miR-24 in primary myogenic progenitors isolated from neither adult nor old mice (**B**). Scale bars: 200μm. n=4-7. Adult: 6-8 months old; old: 20-24 months old; human: young-adult. Two-tailed unpaired Student t-test compared to control. p-value < 0.05 was considered as statistically significant (* p < .05; ** p < .01; *** p < .001). Error bars show S.E.M.

**Figure S5.** CM-DCFH staining negative control. Scale bars: 200μm.

**Supplementary table 1:** Age and gender of the patients donating a muscle biopsy for the isolation of human primary myogenic progenitor cells.

**Supplementary table 2:** Antibodies used for the isolation of satellite cells by FACS.

**Supplementary table 3:** List of primers and oligos used for the study.

**Supplementary table 4:** Table of reagents used for the experiments.

## Acknowledgements

The authors would like to thank Caroline Rainer (Technology Directorate, MARIAC, University of Liverpool), for her help with the isolation of satellite cells; Dr. Aldorada Pisconti (University of Liverpool) for the anti-α7-integrin gift; Dr. Rachel McCormick (University of Liverpool) for helping with 5’UTR GFP construct and Mr. Andy Molloy and members of the Aintree University Hospital (Liverpool) for providing human samples.

## Author contributions

AS, JG, KW performed the experiments; AS, JG, DB and KW performed data analyses; all authors contributed to experimental design, statistical analyses and manuscript preparation.

