## Supplementary figures and tables for "miR-24:Prdx6 interactions regulate oxidative stress and viability of myogenic progenitors during ageing"

**Fig. S1**

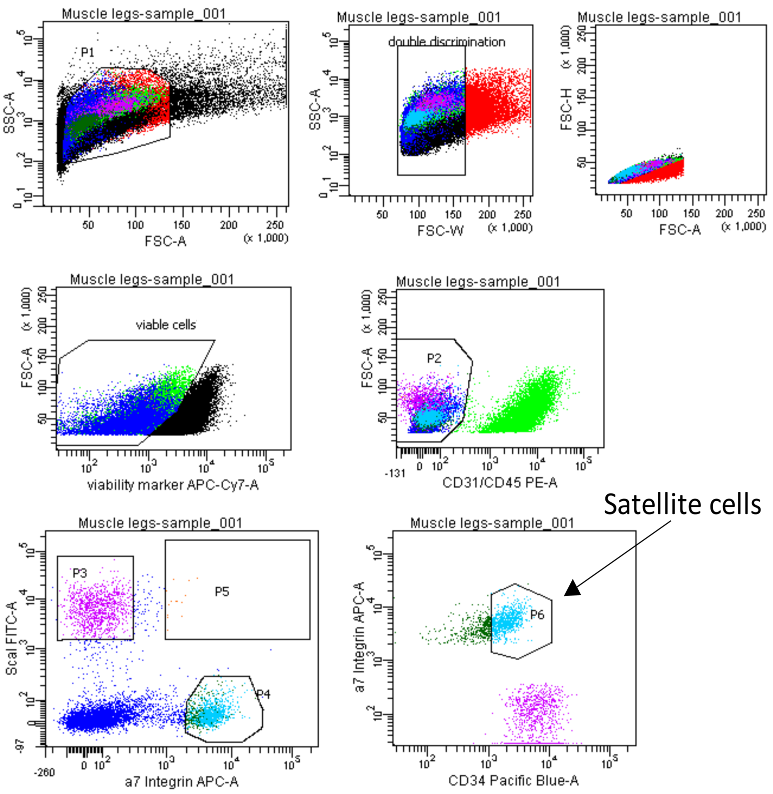
Muscle legs-sample_001
P1
SSC-A
FSC-A
double discrimination
FSC-W
FSC-H
viable cells
viability marker APC-Cy7-A
P2
CD31/CD45 PE-A
P3
P5
P4
ScaI FITC-A
a7 Integrin APC-A
P6
CD34 Pacific Blue-A
Satellite cells

#### Fig. S2

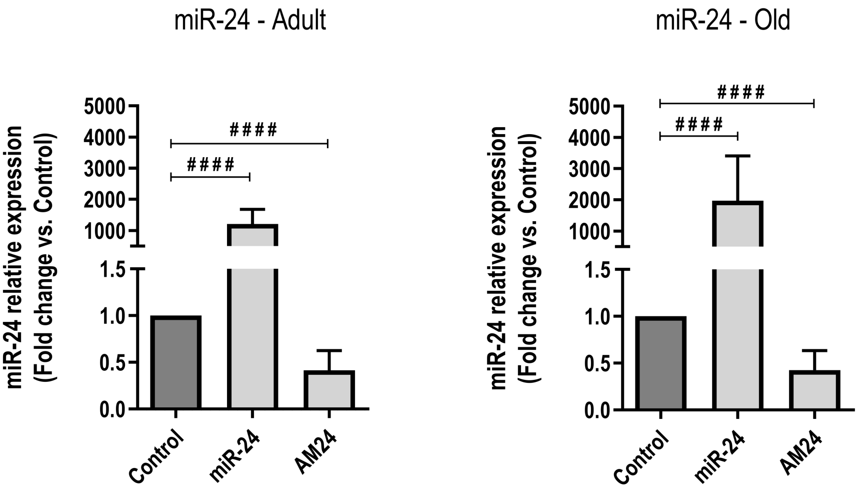
miR-24 - Adult
miR-24 - Old
miR-24 relative expression
(Fold change vs. Control)
5000
4000
3000
2000
1000
1.5
1.0
0.5
0.0
####
Control
miR-24
AM24

#### Fig. S3

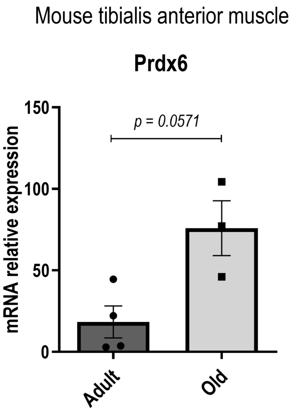
Mouse tibialis anterior muscle
Prdx6
mRNA relative expression
150
100
50
0
p = 0.0571
Adult
Old

### Fig. S4

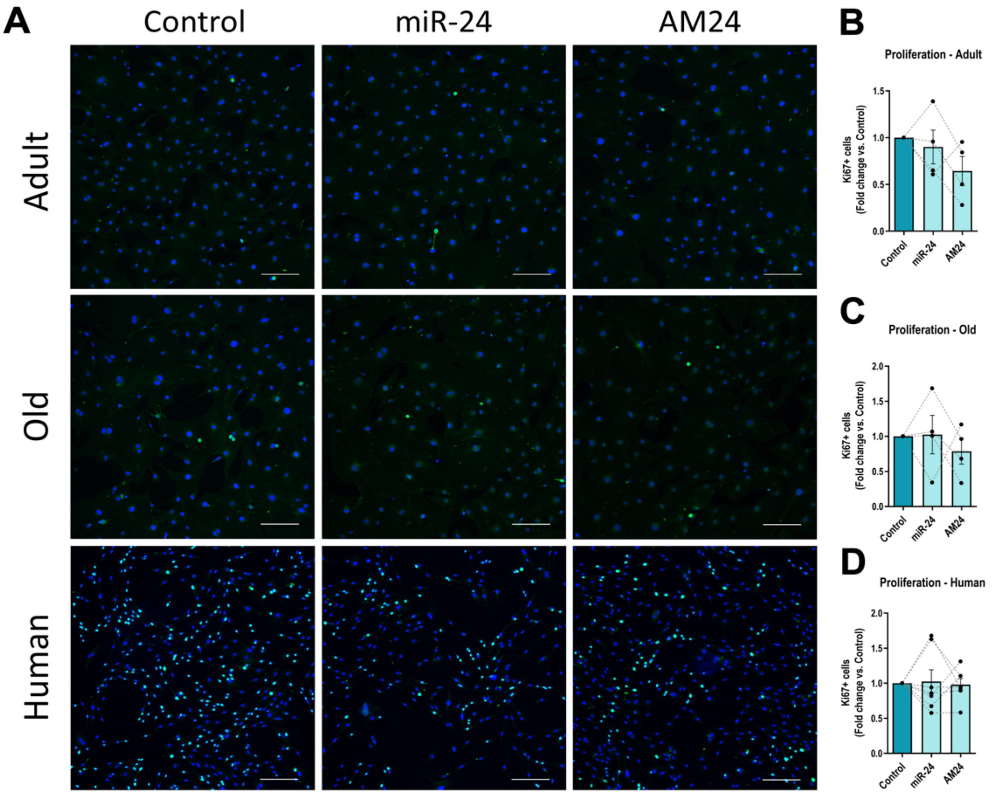


### Fig. S5

CM-$H_2$DCFH staining negative control

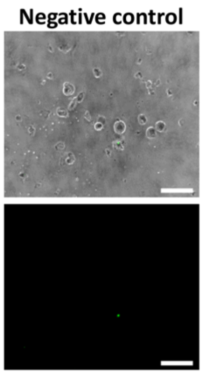


| Donor ID | Age (years) | Gender | BMI |
|---|---|---|---|
| Donor 1 | 32 | Female | <25 |
| Donor 2 | 34 | Female | <25 |
| Donor 3 | 32 | Female | <25 |
| Donor 4 | 31 | Female | <25 |
| Donor 5 | 22 | Female | <25 |
| Donor 6 | 35 | Female | <25 |

**Supplementary table 1:** Age and gender of the patients donating a muscle biopsy for the isolation of human primary myogenic progenitor cells.

| FACS Conjugated antibody | Company | Catalogue number | Dilution | FACS Aria filter |
|---|---|---|---|---|
| Anti-CD31-PE: PE Rat Anti-Mouse CD31. Clone: MEC 13.3. Isotype: Rat IgG2a, κ. 0.2 mg/ml | BD Biosciences PharmingenTM | 561073 | 1:1333 | 575/26 (PE) |
| Anti-CD-45/PE: PE Rat Anti-Mouse CD45. Clone: 30-F11. Isotype: Rat IgG2b, κ. 0.2 mg/ml | (BD Biosciences PharmingenTM | 553081 | 1:1333 | 575/26 (PE) |
| Anti-Sca1/FICT: FICT Rat Anti-mouse Ly-6A/E. Clone: E13-161.7. Isotype: Rat IgG2a, κ. 0.5 mg/ml | BD Biosciences PharmingenTM | 553335 | 1:1333 | 530/30 (FITC) |
| Anti-Alpha 7 Integrin 647. Clone: R2F2. Isotype: Rat IgG2b. 1.0 mg/ml | AbLab | N/A | 1:2000 | 660/20 (APC) |
| BV421 Rat Anti-Mouse CD34 Clone RAM34 (RUO). 0.2 mg/ml | BD Biosciences PharmingenTM | 562608 | 1:1000 | 450/40 (Pacific Blue) |
| Fixable Viability Dye eFluor 780 (label dead cells) | Affimetrix eBiosciences | 65-0865-14 | 1:4000 | 780/60 (APC-Cy7) |

**Supplementary table 2:** Antibodies used for the isolation of satellite cells by FACS.

| Gene | Company | Organism | Sequence (5'-3')/ Cat. Number |
|---|---|---|---|
| Hs_SNORD61_11 miScript Primer Assay | Qiagen | Human, mouse | MS00033705 |
| Hs_miR-24_1 miScript Primer Assay | Qiagen | Human, mouse | MS00006552. Targets mature miR: UGGCUCAGUUCAGCAGGAACAG |
| Beta-actin Forward | Sigma-Aldrich | Mouse | GATCAAGATCATTGCTCCTCCTG |
| Beta-actin Reverse | Sigma-Aldrich | Mouse | AGGGTGTAAAACGCAGCTCA |
| 18S rRNA Forward | Sigma-Aldrich | Mouse | CGGCTACCACATCCAAGGAAGG |
| 18S rRNA Reverse | Sigma-Aldrich | Mouse | CCCGCTCCCAAGATCCAACTAC |
| Beta-2 microglobulin Forward | Sigma-Aldrich | Mouse | GGAGAATGGGAAGCCGAACA |
| Beta-2 microglobulin Reverse | Sigma-Aldrich | Mouse | TCTCGATCCCAGTAGACGGT |
| p16 Forward | Sigma-Aldrich | Mouse | TGGTCACTGTGAGGATTCAGC |
| p16 Reverse | Sigma-Aldrich | Mouse | GTTGCCCATCATCATCACCTGG |
| p21 Forward | Sigma-Aldrich | Mouse | ATCCAGACATTCAGAGCCACAG |
| p21 Reverse | Sigma-Aldrich | Mouse | TCGGACATCACCAGGATTGG |

| | | | |
|---|---|---|---|
| Prdx6 Forward | Sigma-Aldrich | Mouse | TTGATGATAAGGGCAGGGAC |
| Prdx6 Reverse | Sigma-Aldrich | Mouse | CTACCATCACGCTCTCTCCC |
| Tumor protein p53 Forward | Sigma-Aldrich | Mouse | CACGTACTCTCCTCCCCTCAAT |
| Tumor protein p53 Reverse | Sigma-Aldrich | Mouse | AACTGCACAGGGCACGTCTT |
| mPRDX6-202 5’UTR Forward | Sigma-Aldrich | Mouse | GCCCCGCCCACTCGGCCAGC |
| mPrdx6-202 5’UTR Reverse | Sigma-Aldrich | Mouse | AGCAACCCTCCGGGCATGGC |
| mPrdx6-202 5’UTR WT | Sigma-Aldrich | Mouse | GCCCCGCCCACTCGGCCAGCACTGATCTAGGTCTCCGCAG**GAGCC**CGCCCGCTGCTCACTGCTGCGGCTGCGCCTCCTTGTTCTCAGCGTCACCACTGCCGCCATGCCCGGAGGGTTGCT |
| mPrdx6-202 5’UTR 24 MUT | Sigma-Aldrich | Mouse | GCCCCGCCCACTCGGCCAGCACTGATCTAGGTCTCCGCAG**GATCC**CGCCCGCTGCTCACTGCTGCGGCTGCGCCTCCTTGTTCTCAGCGTCACCACTGCCGCCATGCCCGGAGGGTTGCT |

**Supplementary table 3:** List of primers and oligos used for the study.

| **Product** | **Company** | **Catalogue number** |
|---|---|---|
| Barium chloride | Sigma-Aldrich | 202738 |
| AllStars Negative Control siRNA Print | Qiagen | 1027280 |
| Syn-mmu-miR-24-3p | Qiagen | MSY0000219 |
| Anti-mmu-miR-24-3p | Qiagen | MIN0000219 |
| Mouse Prdx6 siRNA | ThermoFisher Scientific | s62375 |
| Human Prdx6 siRNA | ThermoFisher Scientific | s18428 |
| Lipofectamine 2000 | ThermoFisher Scientific | 11668019 |
| MF20 primary antibody. Antigen: myosin, sarcomere (MHC). 211 ug/ml | Developmental Studies Hybridoma Bank | MF20-c 2ea |
| Rabbit mAb to Ki67 [SP6]. | Abcam | ab16667 |
| Goat anti-Mouse IgG (H+L) Secondary Antibody, Alexa Fluor 488 conjugate. | Invitrogen | A-11029 |
| Goat anti-Rabbit IgG (H+L) Secondary Antibody, Alexa Fluor 488 conjugate. | ThermoFisher Scientific | A-11034 |
| DAPI (4',6-Diamidino-2-Phenylindole, Dihydrochloride) | Sigma-Aldrich | D9542 |
| Senescence β-Galactosidase Staining Kit | Cell Signaling Technology | 9860 |

| | | |
|---|---|---|
| Acridine Orange hydrochloride solution, 10 mg/mL in $H_2O$ | Sigma-Aldrich | A8097 |
| Ethidium bromide solution. BioReagent, for molecular biology, 10 mg/mL in $H_2O$ | Sigma-Aldrich | E1510 |
| Methanol | Fisher | M/4000/PC17 |
| PBS (immunostaining) | Sigma-Aldrich | P4417 |
| Tween-20 | Sigma-Aldrich | P1379 |
| Wheat Germ Agglutinin (WGA), Fluorescein | Vector Laboratories | FL-1021 |
| Fluoromount | ThermoFisher | 00-4958-02 |
| DAPI (4',6-Diamidino-2-Phenylindole, Dihydrochloride). 1mg/ml | Sigma-Aldrich | D9542 |
| miRNeasy Mini Kit | Qiagen | 217004 |
| TRIzol Reagent | Life Technologies | 15596-018 |
| Chloroform:Isoamyl alcohol 24:1 | Sigma-Aldrich | C0549 |
| Isopropanol | Sigma-Aldrich | I9516 |
| RNAse-free water | Sigma-Aldrich | 3098 |
| Sodium acetate | Sigma-Aldrich | S2889 |
| Nanodrop 2000 | ThermoFisher Scientific | N/A |
| Superscript II Reverse Transcriptase | Life Technologies | 18064 |
| Random Hexamers (50 µM) | ThermoFisher | N8080127 |
| 25X dNTP Mix (100 mM) | ThermoFisher | 4368814 |
| RiboLock RNase Inhibitor (40 U/µL) | ThermoFisher | EO0381 |
| miRScript RT II | Qiagen | 218161 |
| miRScript SybrGreen PCR Kit | Qiagen | 218073 |
| T100 Thermal Cycler | Bio-Rad | 1861096 |
| CFX Connect Real-Time PCR Detection System | Bio-Rad | 1855201 |
| RNU-6 qPCR primer | Qiagen | MS00033740 |
| Snord-61 qPCR primer | Qiagen | MS00033705 |
| miR-24_1 miScript Primer Assay | Qiagen | MS00006552 |
| Select agar | Sigma-Aldrich | A5054 |
| MyTaq Red Mix | Bioline | BIO-25043 |
| GeneJET Genomic DNA Purification Kit | Thermo Scientific | K0721 |
| One Shot TOP10 Chemically Competent E. coli | Invitrogen | C404010 |
| SYBR Safe DNA Gel Stain | Invitrogen | S33102 |
| DNA Gel Loading Dye (6X) | Thermo Scientific | R0611 |
| UltraPure Agarose | Invitrogen | 16500500 |
| GFP Tag Antibody, ABfinity Rabbit Monoclonal | ThermoFisher Scientific | G10362 |

| | | |
|---|---|---|
| CM-H2DCFDA (General Oxidative Stress Indicator) | Invitrogen | C6827 |
| FLUOstar OPTIMA microplate reader | BMG Labtech | N/A |
| Hydrogen peroxide solution 30 % (w/w) in H2O, contains stabilizer | Sigma-Aldrich | H1009 |
| 1x RBC (Red Blood Cell) Lysis Buffer | eBioscience | 00-4333-57 |
| FACS Aria III Flow Cytometer | BD Biosciences | N/A |
| C1 confocal laser scanning microscope system. 10x magnification. Eyepieces: CFI 10x/22. Blue: ex.: 405 nm, em.: 450/35; Green: ex.: 488 nm, em.: 515/30; Red: ex.: 543, em: 605/15 (for WGA, MF20, Live/dead, Ki67 and CM-$H_2$DCFDA) | Nikon | N/A |
| Axiovert 200 inverted microscope. 10x magnification. Eyepieces: Carl Zeiss 1016-758 W-PI 10x/25 (for H&E and senescence-associated β-Galactosidase staining) | Carl Zeiss | N/A |

**Supplementary table 4:** Table of reagents used for the experiments.
